# Deformed Alignment of Super-Resolution Images for Semi-flexible Structures in 3D

**DOI:** 10.1101/461913

**Authors:** Xiaoyu Shi, Galo Garcia, Yina Wang, Jeremy Reiter, Bo Huang

**Affiliations:** Department of Pharmaceutical Chemistry, University of California, San Francisco, San Francisco, CA 94143, USA; Department of Biochemistry and Biophysics, University of California, San Francisco, San Francisco, CA 94143, USA; Cardiovascular Research Institute, University of California, San Francisco, San Francisco, CA 94143, USA; Chan Zuckerberg Biohub, San Francisco, CA 94158 USA

**Keywords:** Super-resolution microscopy, Image analysis, Image correlation, Single-particle image alignment, Single-molecule localization

## Abstract

Due to low labeling efficiency and structural heterogeneity in fluorescence-based single-molecule localization microscopy (SMLM), image alignment and quantitative analysis is often required to make accurate conclusions on the spatial relationships between proteins. Cryo-electron microscopy (EM) image alignment procedures have been applied to average structures taken with super-resolution microscopy. However, unlike cryo-EM, the much larger cellular structures analyzed by super-resolution microscopy are often heterogeneous, resulting in misalignment. And the light-microscopy image library is much smaller, which makes classification not realistic. To overcome these two challenges, we developed a method to deform semi-flexible ring-shaped structures and then align the 3D structures without classification. These algorithms can register semi-flexible structures with an accuracy of several nanometers in short computation time and with greatly reduced memory requirements. We demonstrated our methods by aligning experimental Stochastic Optical Reconstruction Microscopy (STORM) images of ciliary distal appendages and simulated structures. Symmetries, dimensions, and locations of protein complexes in 3D are revealed by the alignment and averaging for heterogeneous, tilted, and under-labeled structures.

## Introduction

In the past decade, the development of localization-based super-resolution microscopy has brought the light microscopy to nanometer scales. Imaging beyond the diffraction limit addressed structural and functional biomedical questions of subcellular organelles that could not be resolved by conventional light microcopy. For example, the *in situ* dissection of macromolecular protein complexes includes ciliary transition zone ^1-3^, neuronal synapses ^4^, nuclear pore complex ^5,6^, focal adhesion complex ^7^, clathrin-coated pits ^8^, centrosome ^9,10^, and the escort complex at viral budding sites ^11^.

To obtain optical super-resolution images, the target proteins or DNA/RNA sequences are fluorescently labeled by organic dye or fluorescent protein tags. However, the targets are usually under labeled. The cause of the low labeling efficiency includes low affinity antibodies, dye quenching, immature fluorescent proteins, and less-than-one dye to antibody conjugation ratios, etc. In certain cases, it might not be possible to achieve full labeling experimentally. Consequently, image alignment and averaging are often required to make accurate conclusions about the biological system of interest. Template-free cryo-EM image alignment procedures have been applied and adapted to average structures taken with super-resolution microscopy in 2D ^12, 13, 14^, and in 3D ^15^. Unlike cryo-EM, the much larger cellular structures analyzed by super-resolution microscopy are often not completely rigid and homogenous. For instance, the diameter of the rings of ciliary distal appendages varies from 369 to 494 nm ^1^. And the shape for the rings are often elliptical due to the flexibility of cilia. In general, many cellular organelles larger than 200 nm are semi-flexible. These semi-flexible structures keep the common symmetry and arrangement pattern, but have heterogeneous size and shape, such as centrioles ^9^, ciliary transition zones, ciliary distal appendages ^1-3^, and pre- and post-synapse ^4^. In this case, direct alignment and averaging will lose structural details. Cryo-EM particle alignment deals with structural heterogeneity using classification and class-averaging ^16-19^. However, super-resolution structure analysis often has much fewer images to start with for the following two reasons. First, there are only a few target organelles in a cell. For instance, usually one embryonic fibroblast cell only develops one primary cilia^1^, and each cell only has two centrioles^9^. Second, different from EM, super-resolution light microscopes detect fluorescence labels instead of electron density. Many structures seriously under-labeled cannot be used for the alignment and averaging. The small number of usable super-resolution images is often not enough to meet the minimal amount required for classification.

Here, we take structural flexibility as one degree of freedom for image registration. We designed algorithms to first deform the semi-flexible structures to fairly uniform shape and size, and then to align the deformed structures based on cross correlations without classification. We specifically developed our algorithm for the ciliary transition zone and distal appendages^1^. This procedure could be generalized for any ring structures, such as nuclear pore complexes, and centrioles.

## Results

### Overview of the workflow

We designed two algorithms to align and average semi-flexible ring structures, as well as flat structures randomly oriented in 3D. The deformed alignment algorithm (Figure 1, 2) is suitable for heterogeneous ring-shaped structures in which the heterogeneity is mainly caused by the flexibility of the structure, for example, transition zone and centrioles. The algorithm first deforms individual structures by circulating the structure, and then aligns the images. The 3D alignment algorithm (Figure 3) was designed for randomly oriented flat structures in 3D with heterogeneity in the *x*-*y* projection. This algorithm rotates the structure in 3D, and align the projection on the *x*-*y* plane. Both deformed alignment algorithm and 3D alignment algorithm share the same alignment metric, the cross correlation with the reference in the frequency domain. And the initial reference is just the average image of the structures to be aligned. After iterations of alignment, all the images in the frequency domain are inverse Fourier transformed to the space domain. The final alignment information is used to deform, translate and rotate the coordinates in the molecule lists of the super resolution images. Both deformed alignment algorithm and 3D alignment algorithm achieved sub-pixel precision and fast alignment for linear computing time.

**Figure 1.**
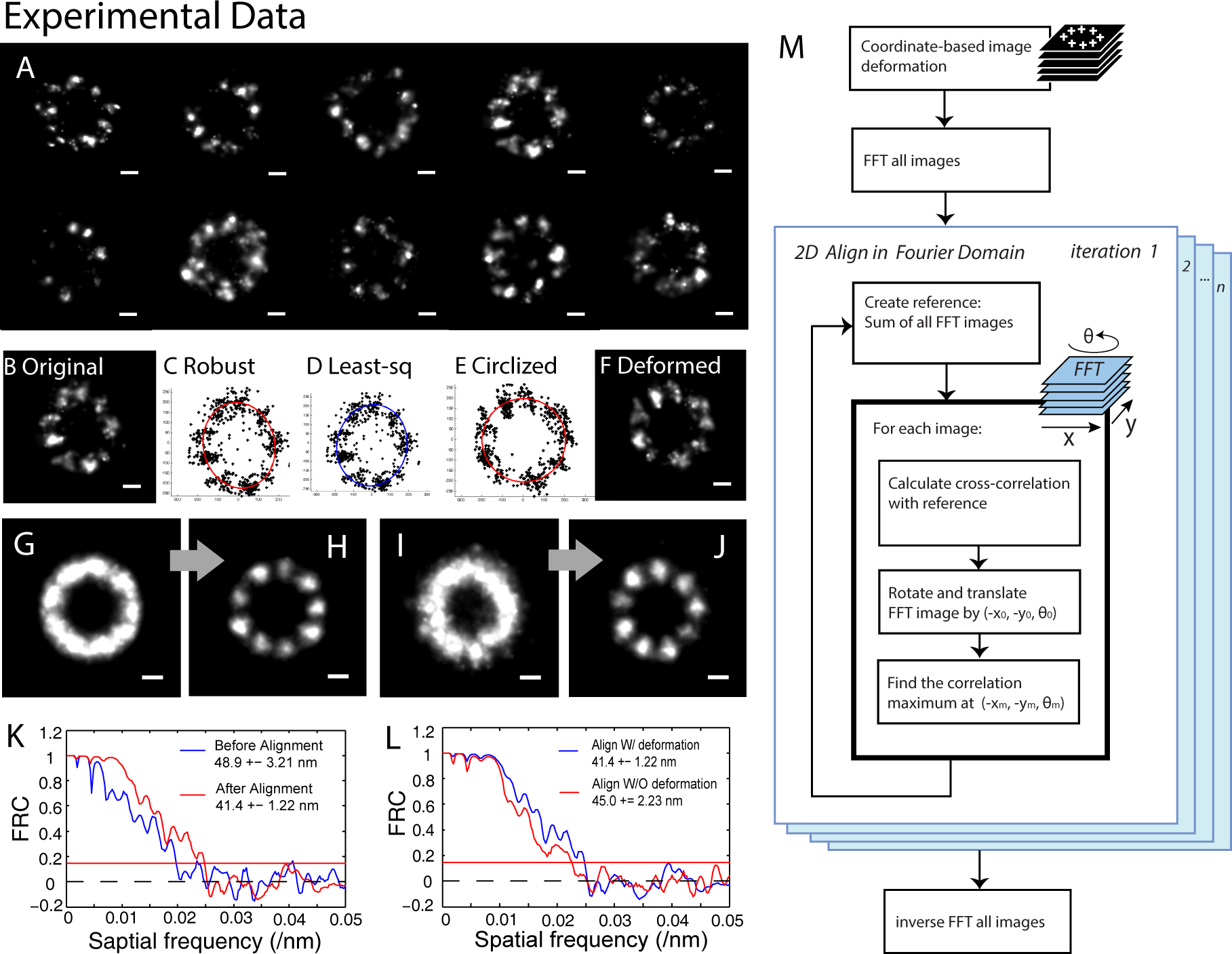
The deformed alignment is demonstrated on real 2D STORM data of the ciliary distal appendages of mTECs cells with CEP164 labeled (A). The STORM localizations of individual structure (B) are fitted to an ellipse. The Robust fitting (C, red) is better than the least-square fitting (D, blue). The ellipse is then deformed to a circle. The coordinates in the structure are moved according to the ellipse-to-circle deformation (E). The circulated structure (F) is normalized to a ring with the fixed diameter. The fixed diameter is the average diameter of total 31 original structures. Images (G) and (H) are the average image of the 31 deformed structures, and their alignment result, respectively. As a comparison, Images (I) and (J) are the average image of the 31 original structures and their alignment result, respectively. Plot (K) includes the FRC curves of average image of original images (blue) and the deformed alignment result (red). Plot (L) compares the FRC curves of the alignment results with deformation (blue) and without deformation (red). The workflow (M) describes how the iterative FFT cross-correlation alignment algorithm aligns super-resolution images in 2D. Scale bar, 100 nm.

**Figure 2.**
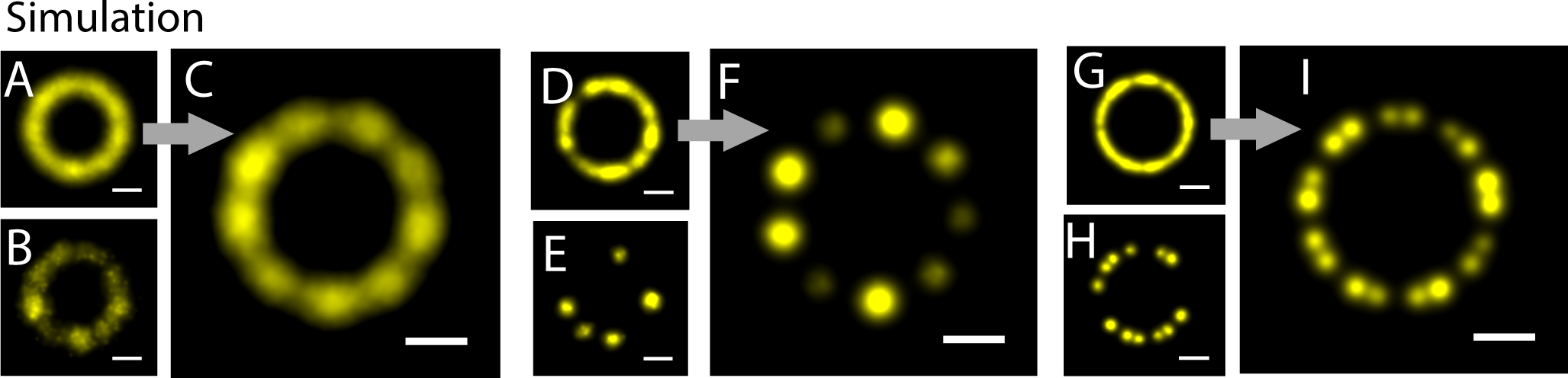
To test the capability of alignment of various localization precision, labeling efficiency, and structural complexity we used simulated rings. Image (A) is the average image of 20 simulated 9-cluster ring structures with the localization precision of 60 nm, and image (B) is one representative of the 20 structures. Image (C) is the deformed alignment result of (A). Image (D) is the average image of 20 simulated 9-cluster rings with 5 cluster labeled at the localization precision of 15 nm, and image (E) is one representative of the 20 structures. Image (F) is the deformed alignment result of (D). Image (G) is the average image of 20 simulated rings with 18 clusters which have alternative 15° and 25° angular spacing between the neighboring clusters at the localization precision of 15 nm, and image (H) is one representative of the 20 structures. Image (I) is the deformed alignment result of (G). Scale bars 100nm.

**Figure 3.**
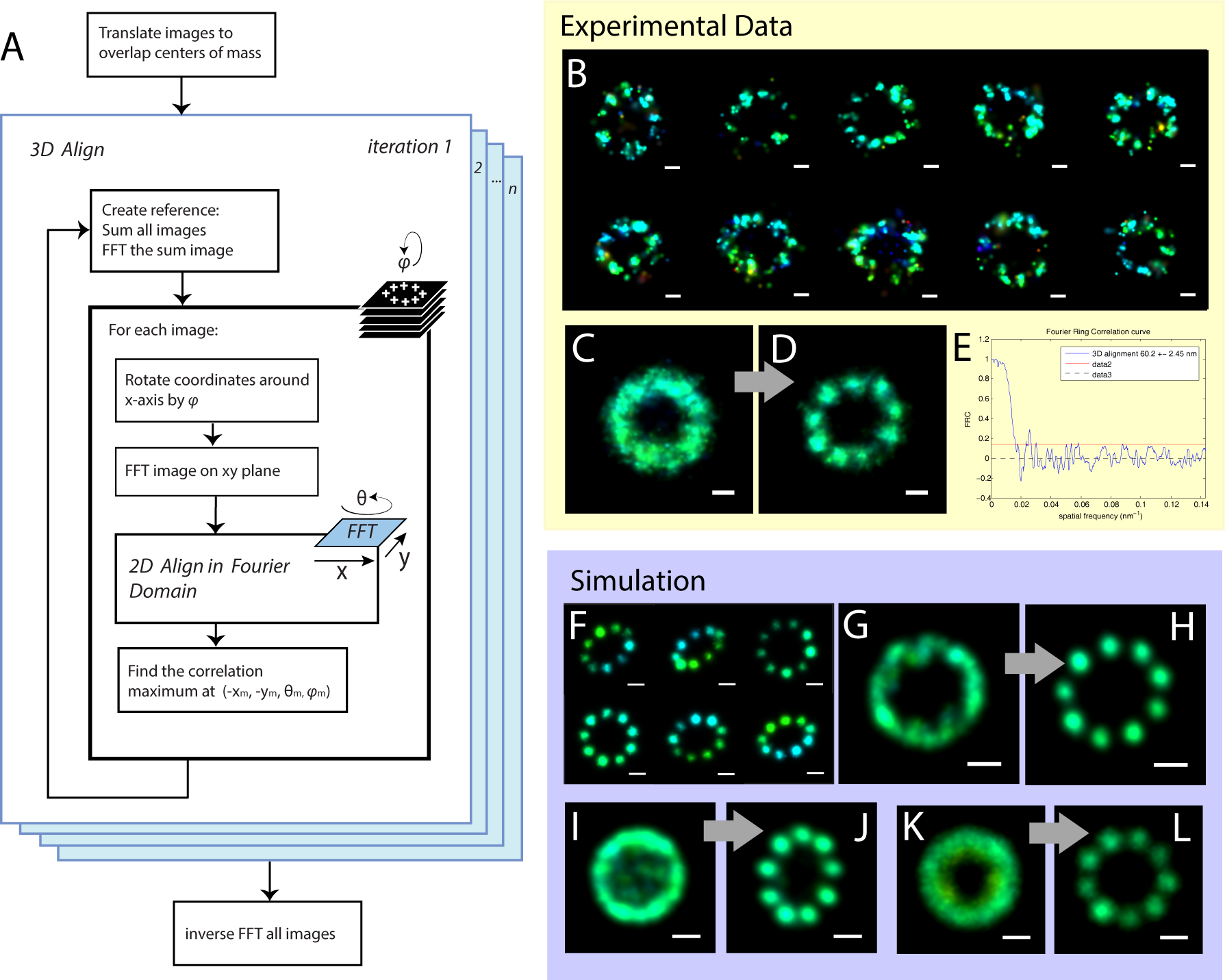
The workflow (A) describes how the iterative cross-correlation alignment algorithm aligns super-resolution images in 3D. The alignment is demonstrated on real 3D STORM data of the ciliary distal appendages of mouse tracheal epithelial cells with CEP164 labeled (B). The average (C) of 29 original individual CEP164 images shows no information about the symmetry of the structure. The average of aligned images (D) shows 9-fold symmetry of the distal appendages labeled by CEP164. (E) FRC analysis of the aligned image in (D). To test the capability of alignment of tilted structures, we used simulated rings with 9 clusters (F). These simulated structures were randomly rotated around x axis within 60°, around z axis within 360°, with a localization precision of 15 nm. Image (G) is the average of 20 original simulated tilted images. Image (H) is the 3D alignment result of (G). Image (I) is the average of 20 simulated structures randomly rotated around x axis within 90°, around z axis within 360°, with a localization precision of 15 nm. (J) is the 3D alignment result of (I). Image (K) is the average of twenty structures simulated localization precision of 60 nm, and randomly rotated around x axis within 30°, around z axis within 360°. (L) is the 3D alignment result of (K). Scale bars 100nm, The localization points are colored by their z coordinate values.

On top of the single-color alignment algorithms, we facilitated multicolor alignment by applying the 3D alignment parameters for the reference channel to other imaged channels (Figure 4). All software codes and sample data in this manuscript are available at https://github.com/Huanglab-ucsf/Deformed-Alignment.

**Figure 4.**
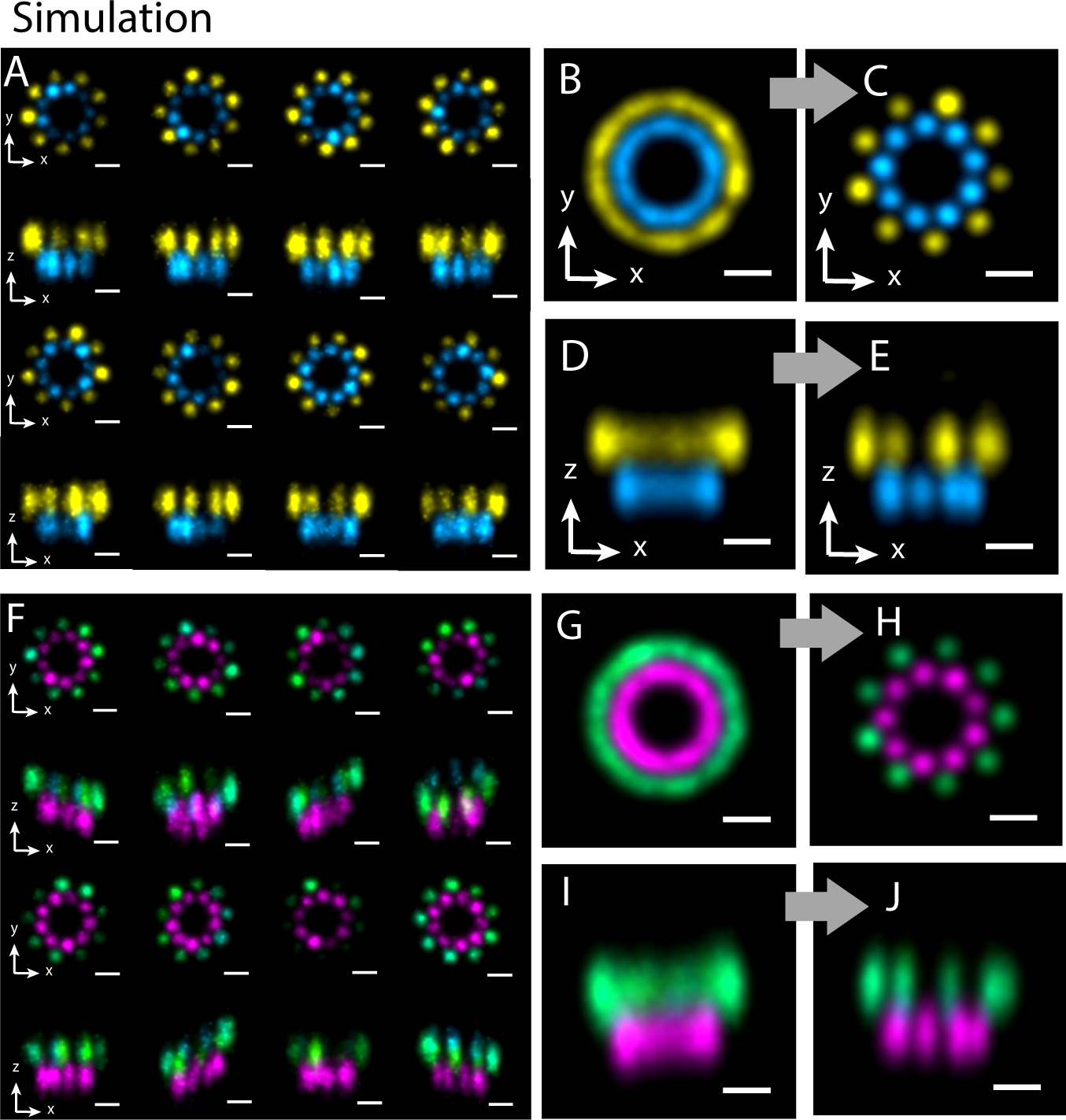
Two-color 3D alignment of simulated structures. (A) Projections of simulated two-color super-resolution images on x-y and x-z planes, with 15 nm lateral localization precision and 30 nm axial localization precision. Image (B) and (D) are the average of 20 original simulated structures on x-y and x-z planes, respectively. Image (C) and (E) are the two-color alignment results on x-y and x-z planes, respectively. (F) Simulated two-color super-resolution images randomly titled around x-axis within 45°, with 15 nm lateral localization precision and 30 nm axial localization precision. (G) and (I) are the average of 20 original tilted simulated structures on x-y and x-z planes, respectively. Image (H) and (J) are the 3D two-color alignment results on x-y and x-z planes, respectively. Scale bars, 100 nm.

### Coordinate-Based Image Deformation

Taking structural flexibility as one degree of freedom for image registration, we deform the semi-flexible structure before aligning the images. Compared to pixel-based images, it is much easier to apply complex deformation functions to coordinate-based images. Fortunately, the data format of localization microscopy is a list of coordinates. Each coordinate (*x*, *y*, *z*) for a 3D image or (*x*, *y*) for a 2D image locates a fluorophore. First, we used robust fitting to fit the coordinates of individual structures to ellipses described as

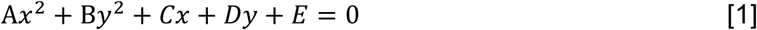

where the algebraic parameters *A* to *E* are converted to geometric parameters center (*x*_0_, *y*_0_), long axis *a*, short axis *b*, and the rotation angle *ϕ*.

Robust fitting is an alternative to least squares fitting when data are contaminated with outliers. We demonstrated that robust fitting gave more accurate results for localization super resolution images, by comparing the ellipse fittings of a STORM image of ciliary distal appendages with robust fitting (Fig 1C) and with least square fitting (Fig 1D), respectively. In this algorithm, coordinates out of 1.5 standard deviations are removed to clean the images. With the geometric parameters obtained from the ellipse fitting, we center the structure to (*x*_0_, *y*_0_) and rotate all the coordinates by – *ϕ* around the center. Finally, we circulate the structure by simply scaling (*x*, *y*) with Eqs. [2] and [3],

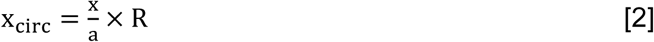

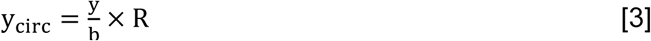

where *a* is the long axis, *b* is the short axis, and *R* is the average radius of all structures to be aligned (Fig 1A).

With experimental data, we demonstrated the improvement of resolution and symmetry in the average images aligned after deformation (Fig 1H), compared to the result of alignment without deformation (Fig 1J). The Fourier ring correlation (FRC) analysis quantified the resolution improvement made by the coordinate-based image deformation (Figs 1K & 1L). The FRC of the average image aligned with deformation is 41.4 ±1.2 nm, while the FRC without the deformation step is 45 ± 2.2 nm.

### Fast Image Alignment using Correlation in Fourier Domain

Using either the deformed or 3D alignment algorithms, 30 images can be aligned in a few minutes with a CPU at 2.8 GHz and 24 GB memory. And the computation time is linear to the number of the images to be aligned. The high-speed image registration is enabled by Discrete Fourier Transform (DFT). The DFT algorithm is a modification of the efficient subpixel image registration algorithm written by Guizar-Sicairos *et al.*^19^ that achieved efficient subpixel image registration by up-sampled DFT cross-correlation. This algorithm registers images using 2D rigid translation. We considered sample rotation and implemented rotational registration (Figure 1M).

Briefly, the original algorithm first obtains an initial 2D shift estimate of the cross-correlation peak by fast Fourier transforming (FFT) an image to register to a reference image. Second, it refines the shift estimation by up-sampling the DFT only in a small neighborhood of that estimate by means of a matrix-multiply DFT. We added a loop to rotate the images to align from 0 to 359° by 1° at a step, and then performing the original first step after each rotation step to obtain 360 cross-correlation peaks. The biggest peak provides not only the initial 2D shift estimate but also the optimal angle of rotation. The images of single structure were aligned for ten iterations. To eliminate the bias that can be introduced by a template, we used the average of all the images to be aligned as the reference image for the first iteration. Each image was translated and rotated to the reference image to maximize the cross-correlation between the image to be aligned and the reference image in the Fourier domain. The average of the images aligned in one iteration was then served as the reference image for the next iteration. Our algorithm aligned the images with a precision of 1/100 of a pixel. The up-sampling rate can be adjusted higher without increasing computation time.

We quantify the alignment using normalized root-mean-square error (NRMSE) *E* between f(x,y) and rotated *g*(*x*, *y*), defined by [4]

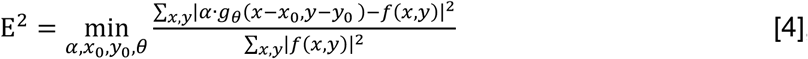

where summations are taken over all image points (*x*,*y*), *α* is an arbitrary constant, *g_θ_*(*x*, *y*) is the image *g*(*x*, *y*) rotated by angle θ_0_ as in equation [5]

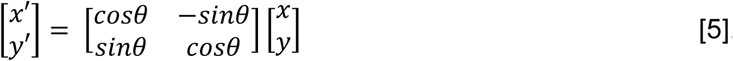

Finding the *x*_0_, *y*_0_, and θ_0_ for the minimum NRMSE is equivalent for the maximum cross-correlation *r*_fg_, defined by [6]

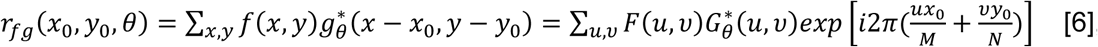

where *N* and *M* are the image dimensions, (*) denotes complex conjugation, *F*(*μ*, *υ*) and *G_θ_* (*μ*,*υ*) are the DFT of *f*(*x*, *y*) and *g_θ_*(*x*, *y*), respectively, as given by the relation [7]

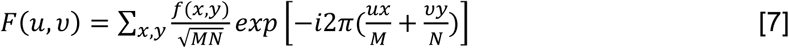

The algorithm was demonstrated with the experimental 2D STORM data of ciliary distal appendages of mouse tracheal epithelial cells (MTECs) with CEP164 labeled by Alexa Fluor 647. The deformed 2D alignment result (Fig 1H) of 31 under-labeled structures with semi-flexible shape and size (Fig 1A) showed clear 9-fold symmetry. The elongated distribution of CEP164 in each distal appendage was also represented in individual subunit of the average image. The elongated distribution of CEP164 in each subunit agrees with the fiber shape of the distal appendage imaged by EM^20^.

To test the capability of our algorithm to align of images with various localization precision, labeling efficiency, and high structural complexity, we simulated three sets of localization images. (1) With the localization precision of 60 nm, we simulated twenty rings, each of which consists of 9 evenly distributed clusters with the diameter of 300 nm (Fig 2A average structure, B single structure). The algorithm was able to resolve 9 clusters clearly by aligning simulated structures with 10 integrations in 363 seconds on a server with 2.8 GHz CPU and 24GB memory (Fig 2C). (2) To test the alignment of structures with low labeling efficiency, we simulated 20 9-cluster rings with 5 random clusters labeled at the localization precision of 15 nm (Fig 2D average structure, E single structure). The average image of aligned underlabled structures recovered clear 9 clusters (Fig 2F). (3) We also evaluated the alignment accuracy with simulated structures with higher structural complexity. We simulated 20 rings with 18 clusters which have alternative 15° and 25° angular spacing between the neighboring clusters (Fig 2G average structure, H single structure). The simulated labeling efficiency is 67%. The average image of aligned structures successfully represented the complex symmetry (Fig 2I).

### 3D Alignment

Based on the 2D alignment algorithm, we developed a 3D alignment algorithm finding the minimum NRMSE (maximum cross-correlation) while rotating and translating the structure in 3D. Any rotation around *y* axis is equal to the combination of rotation around *x* axis and 90 rotation around *z*. So, the 3D alignment algorithm (Fig 3A) adds only one for-loop out of the 2D alignment algorithm described above and excludes the deformation step. This for-loop rotates individual structures around its x axis. The localization coordinates (*x*, *y*, *z*) are rotated by angle *ϕ* in a preset range. The coordinates are transformed by operation

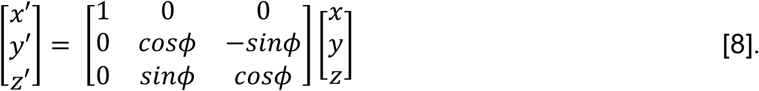

The workflow is described in detail in Fig 3A. The structures are aligned finding the *x*_0_, *y*_0_, *θ*_0_, and *ϕ*_0_ for minimum NRMSE or maximum cross-correlation *r*_fg_.

We used the algorithm to align 29 experimental 3D STORM images of ciliary distal appendages with long axes varying from 376 to 470 nm (Fig 3B). These *in situ* structures were randomly oriented in 3D, and their labeling efficiency was about 60-85%. Before alignment, the average structure showed a ring shape but with no information on the angular distribution of the clusters. After 5 iterations of 3D alignment, the average image showed 9 clusters with approximately even angular distribution around the ring.

To test the capability of alignment of largely tilted structures, we simulated ten rings with 9 clusters. These simulated structures are randomly rotated around the x axis within 60° and around the z axis within 360°, at a localization precision of 15 nm (Fig 3F). The average of simulated ring images before alignment shows no information about the symmetry of the structure (Fig 3G). The 3D alignment result shows 9-fold symmetry (Fig 3H). The algorithm also provides good alignment results for simulated structures with a large tilting angle (90°) around x-axis (Fig 3I, J), and for large localization precision (60 nm) in 3D (Fig 3K, L). These results indicate that our 3D alignment algorithm can accurately register largely titled and noisy images, and can efficiently extract the common structural pattern among these heterogeneous images.

### Two-Color Alignment

The two-color alignment includes two steps. We first picked the channel with higher resolution as the reference channel and aligned this channel with the algorithm described above. Second, the alignment parameters for the reference channel were applied to the other channel.

To test the algorithm of two-color alignment, we simulated twenty two-color super-resolution images with 15 nm lateral localization precision and 30 nm axial localization precision (Figure 4A). Each simulated structure is composed of two parallel rings with diameters of 200 and 300 nm, respectively. Each ring has 9 clusters that are evenly distributed. The average number of localizations in each cluster is 50. The angular distributions of the clusters of the smaller rings and the bigger ring are offset by 20°. The two rings are 100 nm distal to each other. Our alignment algorithm successfully aligned the images. The aligned average image shows 9-fold symmetry and the 20° angular offset between the two rings in the top view (Fig 4C), and 100 nm distal distance between the two rings in the side view (Fig 4E).

To validate the algorithm’s capability of aligning 2C images that are tilted in 3D, we randomly rotated the structures used in the case above around x axis in a range of 45°, and around z axis in a range of 360° (Fig 4F). The 3D alignment algorithm again efficiently aligned the titled structures and recovered the 3D arrangement in the average structure (Fig 4H and J).

### Conclusion

We have demonstrated a deformed 2D alignment algorithm that can accurately align semi-flexible ring structures from both experimental STORM images of ciliary distal appendages and simulated images with high noise and complexity. Information on symmetry and common structural features were efficiently extracted from a few tens of heterogeneous structures. The cross-correlation based alignment algorithm is largely accelerated by DFT and registration in the frequency domain. We also demonstrated that our 3D alignment algorithm can accurately align images of structures tilted in space, using 3D STORM images of ciliary distal appendages, and simulated structures randomly rotated around *x* axis. For two-color alignment, we simply applied the alignment parameters for the reference channel to the other channel, and achieved sub-pixel alignment of two-color super-resolution images. The two-color algorithm can be easily expanded to multiple color alignment. Use of these algorithms makes accurate registration of super-resolution images within a hundredth of a pixel. Information on the general geometric feature of heterogeneous structures can be extracted with tens of images. All the algorithms including deformed 2D, 3D, and two-color alignments are computationally manageable on a regular desktop computer. The deformed 2D algorithm can be applied to any semi-flexible ring-shaped structures. The 3D and multicolor algorithms will provide substantial advantage for any applications that require aligning and averaging super-resolution images in 3D.

## Acknowledgements

This work is invoked by Dr. Joerg Schnitzbauer *et al.*’s Correlation analysis framework for localization-based super-resolution microscopy. We are grateful for Dr. Joerg Schnitzbauer for sharing the code. We thank Drs. David Agard, Yifan Cheng, Shenping Wu and Jean Paul Armache for their consultation on the EM particle alignment and averaging. This work was also highly inspired by personal conversations with Juan Guan, Dan Xie, and Bin Yang, and discussions with all the other Huang lab members. This project is supported by the National Institutes of Health (Director’s New Innovator Award DP2OD008479 and R01GM124334 to B.H., R01AR054396 and R01GM095941 to J.F.R., F32GM109714 to G.G.III), and by the NIH Pathway to Independence Award (1K99GM126136) and the UCSF Mary Anne Koda-Kimble Seed Award for Innovation to X.S.. B.H. is a Chan Zuckerberg Biohub investigator.

